# Intrinsic protein disorder reveals divergent signaling architectures and lifestyle adaptation in *Entamoeba* species

**DOI:** 10.64898/2026.08.12.744381

**Authors:** Rooksana E. Noorai, Stevin Wilson, Lesly A. Temesvari

## Abstract

Intrinsic protein disorder plays a central role in cellular regulation and host-pathogen interactions, yet its proteome-wide distribution and functional organization in amoebozoan parasites remain poorly understood. Here, we present the first comparative analysis of intrinsic disorder across the proteomes of *Entamoeba histolytica*, which causes amoebic dysentery, and two related species, *E. dispar*, and *E. invadens*. For comparison, we analyzed intrinsic protein disorder in two phylogenetically distant parasites, *Plasmodium falciparum* and *Trypanosoma brucei*. Overall, *Entamoeba* species exhibited significantly lower levels of intrinsic disorder than *P. falciparum* and *T. brucei*. Despite this, all organisms displayed a conserved functional trend in which increasing disorder was associated with a shift from metabolic and catalytic processes toward gene expression-related functions. However, notable organism-specific differences emerged. *P. falciparum* showed persistent enrichment of gene expression functions across all disorder levels, while *T. brucei* maintained metabolic, redox, and transport processes throughout the disorder spectrum. In contrast, *Entamoeba* species uniquely retained GTPase- and phosphorylation-associated signaling across all levels of disorder, with the strongest enrichment observed in the pathogenic species, *E. histolytica* and *E. invadens*. This pattern likely reflects a reliance on rapid environmental sensing, cytoskeletal remodeling, and vesicle trafficking, necessary for successful infection. Consistent with this, there was reduced enrichment of G-protein signaling in the non-pathogenic commensal, *E. dispar*, especially in highly disordered proteins. Secretome analysis further revealed that, unlike *P. falciparum*, *E. histolytica* possesses a more structurally ordered secretome, suggesting selection for stable catalytic proteins in the host intestinal environment. Finally, no consistent relationship was identified between intrinsic disorder and vaccine efficacy for several key antigenic targets. Together, these findings demonstrate that intrinsic disorder is differentially deployed across parasites, and utilization of disordered proteins has diverged in accordance with each organism’s lifecycle and host interaction strategy. These findings highlight the role of intrinsic disorder in shaping parasitism.

## Introduction

Intrinsically disordered proteins (IDPs) are a class of proteins that exist without permanent, well-defined three-dimensional structures in physiological conditions (reviewed in [1]). They are aptly described as “dancing protein clouds” [2] because they exhibit a dynamic architecture that enables functional adaptability. Intrinsic disorder can encompass entire proteins or discrete regions within otherwise structured proteins. These regions are often called intrinsically disordered regions (IDRs). Over the past few decades, the concept of intrinsic disorder has emerged as a novel paradigm in protein biochemistry, reshaping the traditional structure-function relationship and highlighting the importance of conformational flexibility in biological systems. IDPs are now recognized across all domains of life, including viruses [3,4], prokaryotes [4,5], and eukaryotes [1,4].

Intrinsic disorder is typically encoded at the amino acid level [6,7]. IDPs and IDRs are routinely enriched in polar and charged residues and depleted in aromatic and hydrophobic amino acids, the latter of which facilitate stable core formation. Proline is particularly abundant in IDRs and is often cited as the most disorder-promoting amino acid residue because its rigid cyclic moiety (pyrrolidine ring) disrupts secondary structure by preventing the formation of α-helices and β-sheets [8]. Enrichment of disorder-promoting amino acids, such as proline, enables IDPs to mediate diverse processes such as signaling, transcriptional regulation, and protein-protein interactions [1].

In parasitic organisms, IDPs may be particularly advantageous. Successful parasites must rapidly adapt to changing environments, evade host immune responses, and interact with multiple hosts, host components, and host systems. The ability of IDPs to bind multiple partners and adopt context-dependent conformations makes them well suited to support these diverse demands. Thus, it is not surprising that IDPs have been implicated in parasite virulence [9–11].

Despite growing interest in IDPs in parasitology, proteome-wide studies of IDPs have only been undertaken for a limited number of organisms, including a helminth (liver fluke), *Fasciola gigantica* [12], several apicomplexan parasites, *Plasmodium falciparum* [11,13], *Toxoplasma gondii* [14], and *Theileria annulata* [15], and several kinetoplastid parasites, *Trypanosoma brucei, Trypanosoma cruzi*, and *Leishmania* spp. [16]. These parasites, which possess intracellular lifestyles (*P. falciparum*, *T. gondii*, *T. annulata*, *T. cruzi*, *Leishmania* spp.) or extracellular lifestyles (*F. gigantica*, *T. brucei*), have provided important insights into the role of intrinsic disorder in antigenic variation (immune evasion) [10,11] and host cell remodeling [9]. However, there remains a notable paucity of studies examining IDPs in the amoebozoan parasites. Here, we address this gap by providing the first systematic analysis of IDPs and IDRs in amoebozoans of the genus, *Entamoeba*.

*Entamoeba histolytica* is the causative agent of amoebic dysentery and liver abscess (reviewed in [17]). It is highly host-specific, with humans and non-human primates serving as the primary natural hosts [18]. In humans, *E. histolytica* is responsible for 50 million symptomatic infections and 40,000 to 100,000 deaths annually worldwide. In accordance with its characteristic fecal-oral route of infection, *E. histolytica* infections are common in areas with poor sanitation, particularly in Africa, Latin America, and Central and South Asia. Other species of this genus include *E. invadens*, which infects reptiles, and *E. dispar*, which is morphologically similar to *E. histolytica* and occurs as a nonpathogenic commensal of the human colon [19].

Amoebozoans, such as the *Entamoebae*, are evolutionarily distant from kinetoplastids and apicomplexans. Unlike intracellular pathogens, which thrive in the protected, nutrient-rich, albeit confined, environment of a host cell, the amoebozoans are extracellular, experience direct exposure to the host immune system, and possess the need to move about and break down host tissue for food. For example, *E. histolytica* exhibits a simple two-stage life cycle consisting of environmentally stable cysts and invasive amoeboid trophozoites that can colonize the human intestine, secrete hydrolases and lytic peptides to destroy colonic epithelium, and penetrate host tissues to cause extraintestinal infections, such as liver abscess [17]. In contrast, *P. falciparum* undergoes complex intracellular and extracellular life cycle transitions spanning mosquito vectors and human host cells (hepatocytes and erythrocytes) (reviewed in [20]), while *T. brucei* remains extracellular, alternates between tsetse fly vectors and mammalian host tissues, and relies on antigenic variation for immune evasion (reviewed in [21]). These differences in niche, life cycles, transmission, and host interaction strategies suggest that the prevalence and functional roles of intrinsic disorder may vary substantially across these parasites.

Here, we demonstrate that the genus, *Entamoeba*, exhibits significantly lower overall intrinsic disorder per protein and per proteome than either *P. falciparum* or *T. brucei*. Gene Ontology (GO) analyses revealed that across all species analyzed, proteins with lower levels of intrinsic disorder were enriched in GO terms associated with metabolic and catalytic activities. These transitioned to gene expression-related terms as intrinsic disorder increased. GO terms associated with signal transduction were enriched across all levels of intrinsic disorder in the *Entamoebae*, which was not observed for *P. falciparum* or *T. brucei*. Furthermore, unlike the secretome of *P. falciparum*, the secretome of *E. histolytica* was not enriched in IDPs relative to its whole proteome. Finally, our analyses indicate that intrinsic disorder may not serve as a robust predictor of vaccine candidate proteins in *E. histolytica*. Overall, our findings highlight differences in the evolutionary and functional roles of protein disorder in parasitic protists.

## Methods

### Data Source

Predicted proteomes analyzed in this study were obtained from publicly available eukaryotic pathogen databases. Specifically, the proteomes of *E. dispar* (amoebadb.org; v50), *E. histolytica* (amoebadb.org; v47), *E. invadens* (amoebadb.org; v47), *P. falciparum* (plasmodb.org; v52), and *T. brucei gambiense* (tritrypdb.org; v52) were downloaded from the most current curated releases available at the time of analysis. Previously published secretome [22,23], membrane antigen [24], and vaccine target [25,26] datasets were incorporated for comparative analyses. Protein identifiers were cross-referenced with current database accession numbers prior to downstream analyses. Any protein that could not be reliably matched to current database accession numbers was excluded from downstream comparisons.

### IDP Predictions and Exclusion Criteria

Intrinsic disorder predictions were performed using the publicly available tool, Disopred3 [27], which employs machine learning approaches trained on experimentally validated disorder data to distinguish ordered and disordered residues. To increase robustness, additional predictions were carried out using IUPred3 [28], which predicts IDRs based on estimated pairwise interaction energies within amino acid sequences, identifying regions unlikely to form stable tertiary structures and AUCpreD [29], a deep learning-based predictor that utilizes sequence profiles and neural network architectures to identify IDRs with high accuracy. All tools were implemented using default parameters. All IDP prediction software programs were executed on the Clemson University Palmetto Cluster [30]. Disorder predictions were subsequently summarized at both the protein and proteome levels for comparative analyses among taxa and functional groups. All output data generated and analyzed in this study are available in Supplementary File 1.

Proteins were excluded from downstream analysis if they met one or more of the following criteria. First, proteins whose sequences contained ambiguous amino acid residues were excluded, as these residues are incompatible with the computational disorder prediction tools employed. Second, proteins for which one or more prediction tools did not produce output were excluded. The majority of these were typically very short sequences, primarily fewer than 5 amino acids in length, which falls below the reliable operating range of the prediction algorithms. Third, proteins for which the amino acid sequence reconstructed from prediction tool output could not be verified to match the original input sequence were excluded due to parsing inconsistencies.

### Calculation of Amino Acid Frequencies

Analysis of the frequency of amino acids present in disordered regions compared to ordered regions was carried out as described [31]. Briefly, protein sequences were partitioned into ordered and disordered residues based on prediction outputs, and the relative frequency of each amino acid was calculated within each category. Enrichment or depletion of individual amino acids in disordered regions was determined by comparing their relative occurrence in disordered versus ordered segments.

### Functional Annotation

Gene Ontology (GO) annotations, defined by the Gene Ontology Consortium [32,33] were used to characterize the predicted functions of proteins exhibiting various levels of disorder content. GO terms associated with biological processes, molecular functions, and cellular components were compiled from available database annotations and used to identify functional trends stratified by percent disorder. Microsoft^®^ Copilot™ (Microsoft Corp., Redmond, WA) was used as a supplementary tool to assist in identifying patterns and trends in Gene Ontology (GO) enrichment results. All patterns were verified manually.

### Statistical Analyses

Statistical analyses were performed using GraphPad Prism Ver 10.6.1 (GraphPad Software, San Diego, CA, USA). Where appropriate, methods most suitable for datasets exhibiting unequal variances and highly unequal sample sizes were employed. Comparisons between datasets were conducted using Unpaired or Welch’s *t*-tests. To quantify the magnitude of observed differences, multiple effect size metrics were calculated, including Cohen’s *d*, Hedges’ *g*, and Glass’s Δ. These complementary measures were included to account for unequal variances between datasets and to provide standardized estimates of effect magnitude. Data distributions were evaluated for symmetry by calculating statistical skewness (g₁). Descriptive statistics are presented as mean ± standard deviation unless otherwise indicated. Statistical significance was assessed using a threshold of *p*<0.05.

### Use of AI-assisted Writing Tools

Draft text for portions of the manuscript was generated with assistance from Microsoft^®^ Copilot™. All references were independently identified and curated by the authors, without the use of the AI tool for literature searching or bibliography construction. All content was reviewed, substantially revised, fact-checked, and approved by the authors. The authors assume full responsibility for the content, accuracy, and scientific integrity of the final manuscript.

## Results

### Overall disorder content in parasite proteomes

Several different approaches are available to predict protein disorder. Although sequence-based disorder predictors have become reasonably accurate, to ensure a robust analysis of the *Entamoeba* proteomes, we used three different algorithms, Disopred3 [27], IUPred3 [28], and AUCpreD [29]. To make certain that the resulting predictions were not an anomaly of our methods, we also analyzed the proteomes of two other parasites, *P. falciparum* and *T. brucei gambiense*, for which protein disorder has been analyzed previously [13,16]. The complete dataset is presented in Supplemental File 1 and summarized in Table 1.

**Table 1.**
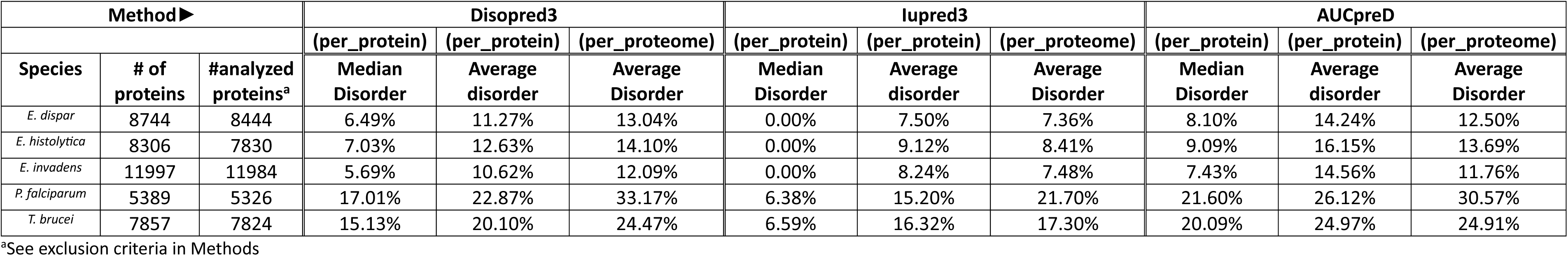
Summary of intrinsic protein disorder parameters in *Entamoeba* spp., *P. falciparum,* and *T. brucei*.

The predictions revealed that *Entamoeba* species possessed lower protein disorder than *P. falciparum* and *T. brucei* (Fig 1). This was the case for both per protein disorder and per proteome disorder. The percentage of per protein and per proteome disorder observed for *P. falciparum* and *T. brucei* was comparable to that observed by others [13,16], which supports the accuracy of our predictions.

**Fig 1.**
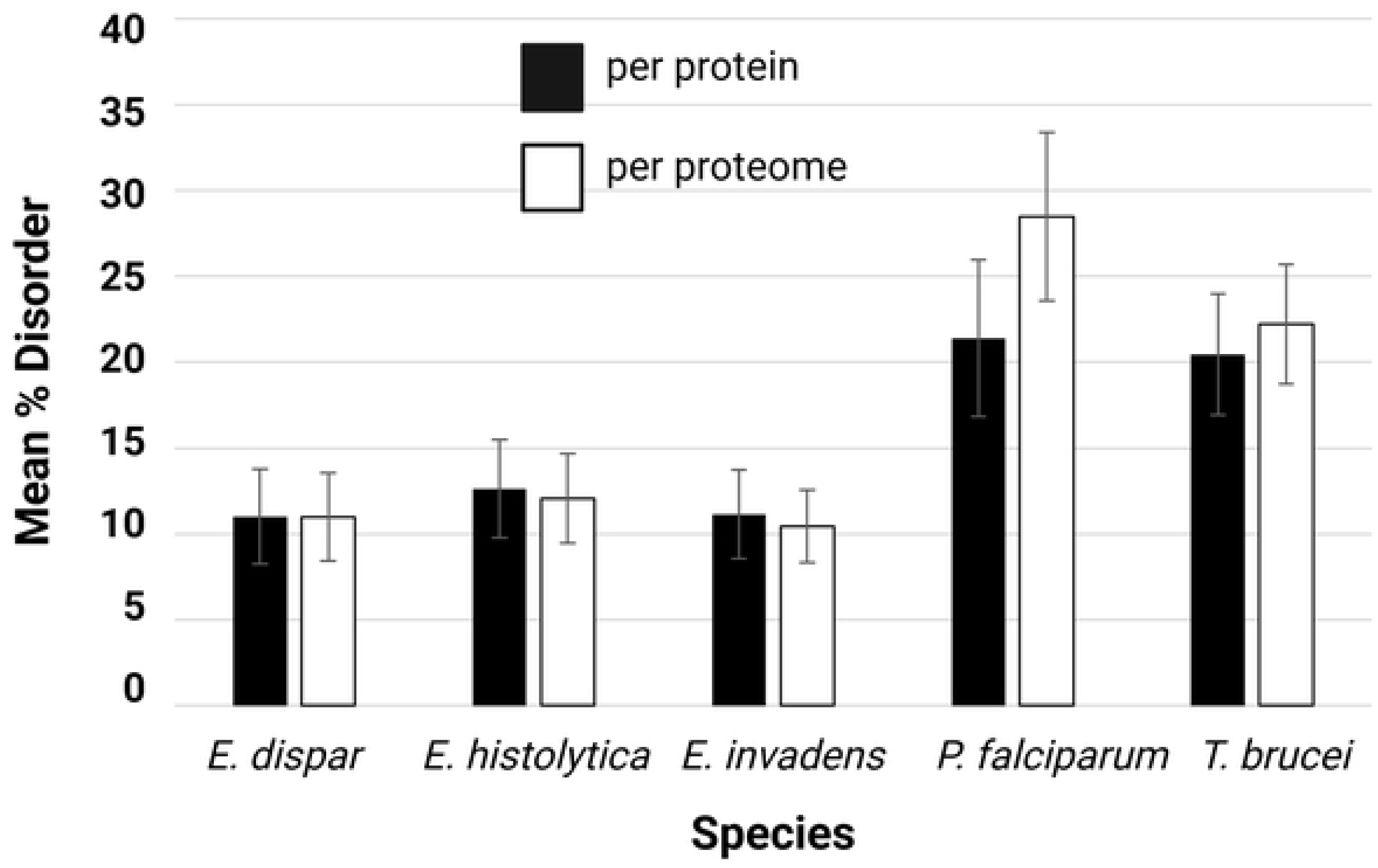
Intrinsic disorder content across selected protozoan species. Mean percentage of intrinsically disordered proteins is shown for *E. dispar*, *E. histolytica*, *E. invadens*, *P. falciparum*, and *T. brucei,* calculated either per protein (black bars) or averaged across the proteome (white bars). Data represent the mean ± SD calculated by three different disorder prediction programs, Disopred3 [27], IUPred3 [28], and AUCpreD [29]. Overall, *P. falciparum* and *T. brucei* exhibit higher levels of disorder compared to *Entamoeba* species.

To statistically compare intrinsic disorder parameters among the parasites, we performed pairwise comparisons using Welch’s *t*-test, which is appropriate because the datasets possess unequal variances and unequal sample sizes. Welch’s *t*-tests demonstrated significant differences among all five groups (all *p*-values < 0.001) (Table 2). *P*-values are strongly influenced by sample size and even negligible differences can appear statistically significant simply due to the power of sample size. Given the large sample sizes of proteomes, we also calculated effect size (Cohen’s *d*, Hedges’ *g*, Glass’s Δ), which assesses the practical significance of our findings, and is less sensitive to sample size. Multiple effect size metrics were used to provide complementary perspectives on group separation. Effect size analyses indicated negligible-to-moderate differences among the *Entamoebae* (Cohen’s *d*, Hedges’ *g*, Glass’s Δ; all<0.78), whereas comparisons between any *Entamoeba* species and either *P. falciparum* or *T. brucei* yielded large effects (Cohen’s *d*, Hedges’ *g*, Glass’s Δ; all ≥1.79). These analyses support the conclusion that *Entamoeba* species possess significantly lower protein disorder than *P. falciparum* or *T. brucei*. The data also demonstrate that intrinsic disorder is not markedly different among the *Entamoebae*.

**Table 2.** Descriptive statistics for mean per protein and per proteome intrinsic disorder among *Entamoeba*, *Plasmodium*, and *Trypanosoma* species.

| *Per Protein* |  |  |  |  |  |  |
| --- | --- | --- | --- | --- | --- | --- |
| Species Comparison<sup>a</sup> | Welch's *t* | *p*-value | Cohen's *d* | Hedges' *g* | Glass's $\Delta$ | Effect Size Interpretation<sup>b</sup> |
| Ed vs Eh | -36.87 | < 0.0001 | -0.58 | -0.58 | -0.57 | Moderate |
| Ed vs Ei | -3.65 | 0.00026 | -0.05 | -0.05 | -0.05 | Negligible |
| Ed vs Pf | -149.48 | < 0.0001 | -2.91 | -2.91 | -2.27 | Extremely large |
| Ed vs Tb | -189.06 | < 0.0001 | -2.99 | -2.99 | -2.67 | Extremely large |
| Eh vs Ei | 37.01 | < 0.0001 | 0.55 | 0.55 | 0.57 | Moderate |
| Eh vs Pf | -124.15 | < 0.0001 | -2.40 | -2.40 | -1.91 | Extremely large |
| Eh vs Tb | -152.00 | < 0.0001 | -2.43 | -2.43 | -2.21 | Extremely large |
| Ei vs Pf | -152.83 | < 0.0001 | -3.07 | -3.07 | -2.24 | Extremely large |
| Ei vs Tb | -200.07 | < 0.0001 | -3.09 | -3.09 | -2.63 | Extremely large |
| Pf vs Tb | 12.63 | < 0.0001 | 0.24 | 0.24 | 0.27 | Small |
| *Per Proteome* |  |  |  |  |  |  |
| Species Comparison | Welch's *t* | *p*-value | Cohen's *d* | Hedges' *g* | Glass's $\Delta$ | Effect Size Interpretation |
| Ed vs Eh | -27.62 | < 0.0001 | -0.43 | -0.43 | -0.42 | Small |
| Ed vs Ei | 14.56 | < 0.0001 | 0.23 | 0.23 | 0.25 | Small |
| Ed vs Pf | -237.27 | < 0.0001 | -4.50 | -4.50 | -3.57 | Extremely large |
| Ed vs Tb | -239.79 | < 0.0001 | -3.87 | -3.87 | -3.22 | Extremely large |
| Eh vs Ei | 44.33 | < 0.0001 | 0.70 | 0.70 | 0.78 | Moderate |
| Eh vs Pf | -209.55 | < 0.0001 | -3.95 | -3.95 | -3.34 | Extremely large |
| Eh vs Tb | -211.42 | < 0.0001 | -3.40 | -3.40 | -2.91 | Extremely large |
| Ei vs Pf | -294.22 | < 0.0001 | -5.24 | -5.24 | -3.67 | Extremely large |
| Ei vs Tb | -293.98 | < 0.0001 | -4.41 | -4.41 | -3.38 | Extremely large |
| Pf vs Tb | 79.29 | < 0.0001 | 1.52 | 1.52 | 1.79 | Large |

<sup>a</sup>Species: *E. dispar* (Ed); *E. histolytica* (Eh); *E. invadens* (Ei); *P. falciparum* 3D7 (Pf); *T. brucei gambiense* (Tb)
<sup>b</sup>Effect size interpretation: negligible (<0.2), small (0.2–0.49), moderate (0.5–0.79), large (0.8–1.19), very large (1.2–1.99), and extremely large (≥2.0). Shaded cells indicate ≥large effect sized (>0.8)

### *Entamoeba* IDRs possess canonical amino acid composition

An analysis of the amino acid frequencies revealed that IDRs in the *Entamoebae* were enriched in polar amino acids such as glutamate (E), lysine (K), glutamine (Q), serine (S), arginine (R), aspartate (D), asparagine (N), and threonine (T) (Fig 2). This was not surprising given that these amino acids are known to be disorder-promoting [6,7]. The cyclic amino acid, proline (P), was also abundant in these regions. Proline hinders the formation of α-helices and β-sheets [8] and frequently participates in protein-protein interactions [34]. Thus, its enrichment in intrinsically disordered protein regions is functionally significant. Like in other systems [6], *Entamoeba* IDRs were depleted in the hydrophobic amino acids, isoleucine (I) and leucine (L), the aromatic amino acids tryptophan (W), tyrosine (Y) and phenylalanine (F), and the thiol-containing amino acid, cysteine (C). Hydrophobic cores of globular proteins are characteristically enriched in I, L, W, Y, and F. Furthermore, C typically contributes stabilizing disulfide bridges to protein regions. Consequently, L, I, F, V, W, Y and C are considered order-promoting residues.

**Fig 2.**
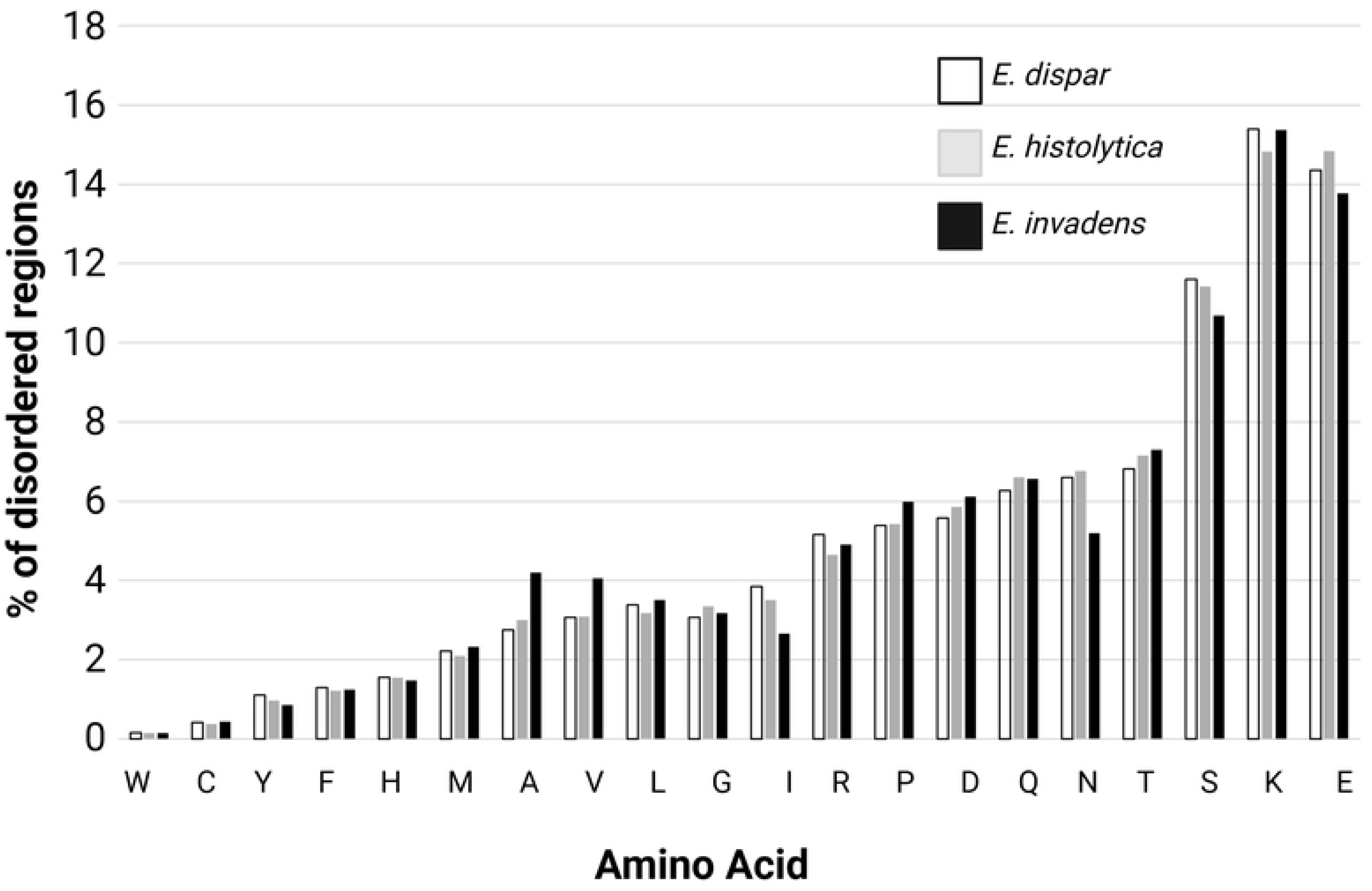
Amino acid composition of intrinsically disordered regions (IDRs) in *Entamoeba* species. Data are presented as the percentage contribution of each amino acid to predicted intrinsically disordered regions in *E. dispar* (white), *E. histolytica* (gray), and *E. invadens* (black). Disorder-promoting residues such as R, P, D, Q, N, T, S, K, and E are enriched across all three species, whereas hydrophobic residues (e.g., W, C, F, Y) are underrepresented. Overall, the three *Entamoeba* species exhibit broadly similar amino acid composition profiles within disordered regions, with minor species-specific variations.

### Distribution of low, moderate, high, and very high disorder proteins across parasite proteomes

Proteome-wide analyses of IDPs often cluster proteins based on the fraction of residues that are predicted to be disordered. For example, proteins can be classified as possessing low disorder (0-10%), moderate disorder (10-30%), high disorder (30-70%) or very high disorder (>70%). Classification of proteins, based on the level of disorder, may provide clues about the types of functions that are likely. For instance, structural proteins often exhibit low disorder [35] while proteins involved in cell signaling, transcription, and chromatin remodeling, which require flexible inter-molecular interactions, often exhibit high disorder (see ref [1]). In the *Entamoebae*, the majority of the proteins (>88%) was asymmetrically distributed in the low (0-10%) or moderate (10-30%) disorder categories (Fig 3). In contrast, in *P. falciparum* and *T. brucei*, a larger fraction of proteins clustered in the high (30-70%) or very high (70%) disorder strata. To quantify these observations, we calculated statistical skewness (*g1*) (Fig 3), which describes the asymmetry or distortion of a distribution. The skewness values were positive for *Entamoeba* species, indicating a long-right tail and clustering on the left (low disorder) whereas the skewness values were negative for *P. falciparum* and *T. brucei* indicating a long-left tail and clustering on the right (high disorder). This supports the conclusion that the *Entamoebae* have lower overall protein disorder than *P. falciparum* and *T. brucei*.

**Fig 3.**
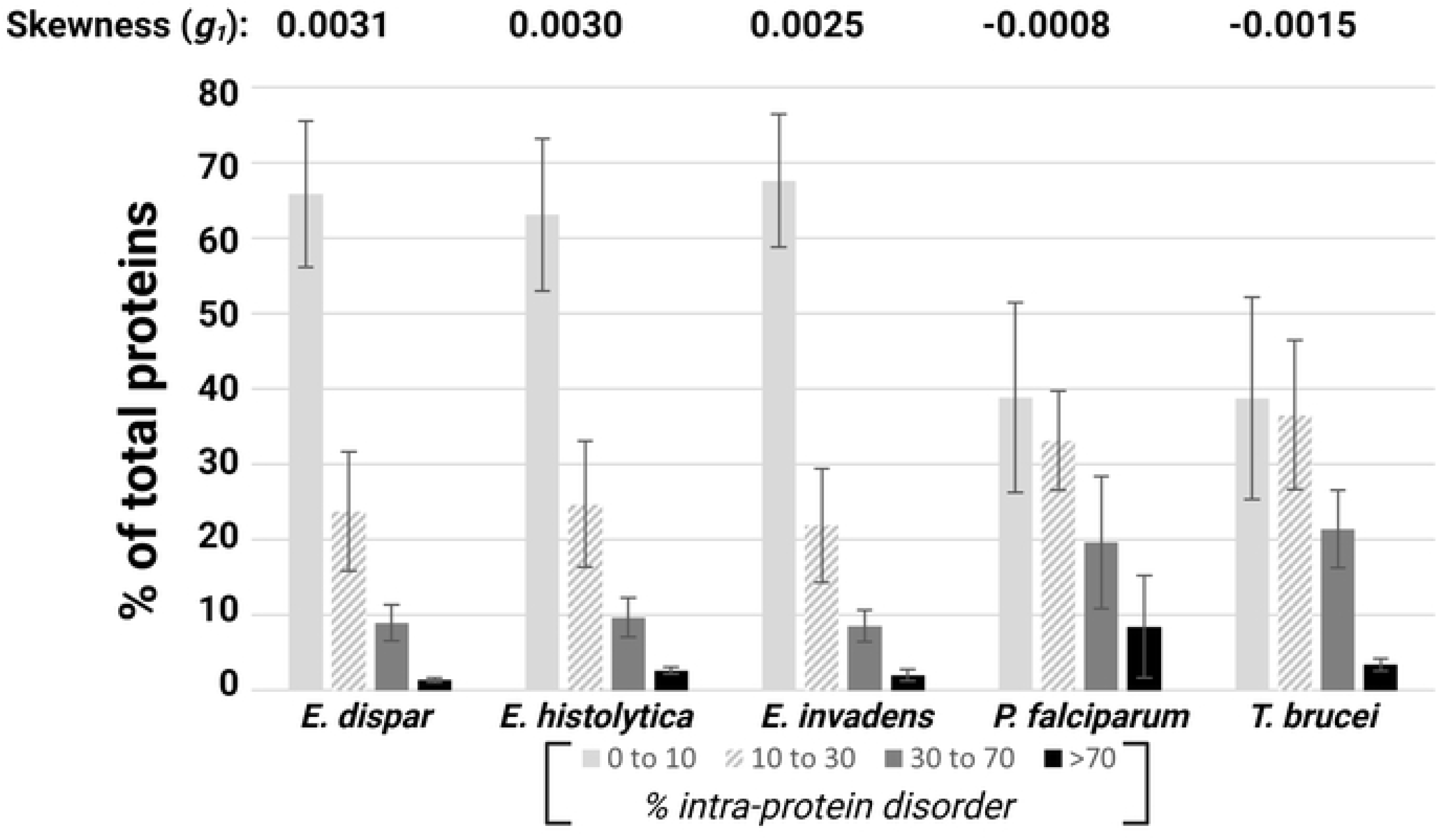
Distribution of proteins by extent of intrinsic disorder across protozoan species. The percentage of proteins within each species, *E. dispar*, *E. histolytica*, *E. invadens*, *P. falciparum*, and *T. brucei* are categorized by the proportion of intra-protein disorder: 0–10% (low disorder; light gray), 10–30% (moderate disorder; hatched), 30–70% (high disorder; dark gray), and >70% (very high disorder; black). The data represent the mean ± SD. Most proteins in *Entamoeba* species fall within the low- and moderate-disorder categories, whereas *P. falciparum* and *T. brucei* display a higher proportion of proteins with moderate to very high levels of disorder. Skewness values (*g₁*) for each species are shown above the plot, indicating a negative skew in *P. falciparum* and *T. brucei* and a positive skew for the *Entamoeba* species.

### Gene ontology (GO) enrichment patterns across intrinsic disorder categories

The relationship between protein structure (including order and disorder) and function prompted us to perform gene ontology (GO) enrichment analyses stratified by low, moderate, high, and very high protein disorder. Proteins with unassigned GO terms, primarily representing hypothetical proteins, represented the most abundant protein class at every level of intrinsic disorder in every species. These unassigned proteins, and the next ten most enriched annotated GO terms (cellular component, molecular function, biological process), along with their percent enrichments, are presented in Supplemental File 2.

GO analysis revealed obvious functional differences across disorder strata. For all five organisms, proteins with low disorder (<10%) were predominantly associated with metabolic and catalytic processes including carbohydrate and lipid metabolism (e.g., GO:0005975, GO:0006629), proteolysis (GO:0006508), and phosphorylation (GO:0006468). However, species-specific trends were evident in this low disorder category. For example, the *Entamoebae* showed marked enrichment in signal transduction and GTPase regulatory activities (e.g., GO:0007165, GO:0005096), whereas *T. brucei* displayed a bias toward oxidoreductase activity (GO:0055114) and transmembrane transport (GO:0055085). *P. falciparum* exhibited enrichment for gene expression-linked terms such as transcription-related processes (GO:0006355, GO:0006351,GO:0000398), translation-related processes (GO:0006412) and ribosomal components (GO:0005840).

At intermediate disorder levels (10-30% and 30-70%), a transitional pattern was observed. The *Entamoebae* retained a strong representation of signaling functions (e.g., GO:0005525, GO:0003924, GO:0005096, GO:0005524, GO:0004672, GO:0004713, GO:0007264, GO:0006468, GO:0043087, GO:0035556, GO:0035023), while *P. falciparum* continued to show enrichment of translation (e.g., G:0006412), transcription (e.g., GO:0006355), and nucleic acid-binding functions (e.g., GO:0003676). *T. brucei* exhibited a broader functional array of enriched GO terms, including cytoskeletal organization (GO:0007018), transport (GO:0055085), and DNA repair processes (GO:0006281), reflecting greater functional diversity in moderately disordered proteins.

Proteins with very high disorder (>70%) converged across all species toward GO terms associated with gene expression and nucleic acid interactions. Enriched categories included translation (GO:0006412), ribosomal structure (GO:0005840), transcription (GO:0006355), RNA binding (GO:0003723), and nucleic acid binding (GO:0003676). Despite this convergence, subtle differences persisted. *P. falciparum* showed the strongest enrichment for transcriptional and RNA processing functions (e.g., GO:0006397, GO:0000398). *T. brucei* retained additional representation of metabolic (GO:0006777), transport (GO:0048208), and cell death-related (GO:0006914) processes. Strikingly, *E. histolytica* and *E. invadens* maintained signaling-associated terms (e.g., GO:0005096, GO:0005525, GO:0007264) even at very high disorder levels. However, in this highest disorder category, the enrichment of signaling-associated terms was less pronounced in *E. dispar* than in the other two *Entamoeba* species.

### AT-richness is not a driver of reduced disorder in the *Entamoebae*

Differences in intrinsic protein disorder could, in principle, be attributed to AT-rich genome composition. AT-richness in DNA is positively correlated with protein disorder because AT-rich sequences tend to encode amino acids that promote protein disorder, such as lysine (K), glutamic acid (E), serine (S), and asparagine (N), while avoiding hydrophobic, order-promoting amino acids. *Entamoeba* species possess high AT genome content (∼70-76.5%, [36]), but this is lower than that found in *P. falciparum* (∼80%, [37]) and higher than that found in *T. brucei* (∼56%, NCBI Assembly GCA_900497135.1 and [38]). Therefore, AT-richness alone cannot explain the observed patterns.

### Structural bias in the *E. histolytica* secretome

*E. histolytica* exhibits a simple two-stage life cycle and occupies only a few hosts (primates) and a few host niches (e.g., intestinal lumen, liver) [17,18]. On the other hand, *P. falciparum* and *T. brucei* exhibit more complex life cycles, with diverse host and vector environments, and more extensive parasite-host interactions in multiple tissue niches. Such complex lifestyles may create a demand for more flexible protein architectures, necessary for binding promiscuity and immune evasion. Thus, higher protein disorder may be an adaptation for host interaction. A prediction of this hypothesis is that disorder should be enriched in proteins that regulate parasite-host interactions.

To test this prediction, we used published secretome datasets from *E. histolytica* [23] and *P. falciparum* [22] and compared the average intrinsic disorder of secreted proteins to that of their respective whole proteomes. Because many parasite-host interactions are mediated through secreted proteins, analysis of the secretomes allowed us to focus specifically on proteins that were most likely to directly interface with the host environment. We were not able to perform a similar analysis for *T. brucei* because a single published secretome dataset [39] could not be reliably reconciled with current database accession numbers, which prevented an unambiguous mapping of proteins between datasets.

Consistent with the prediction, the *P. falciparum* secretome exhibited a significantly higher mean disorder than its corresponding whole proteome (Table 3, Welch’s *t*-test, t ≈ 3.07, p ≈ 0.003). The magnitude of this difference was large, with effect sizes of Cohen’s *d* ≈ 1.90, Hedges’ *g* ≈ 1.88, and Glass’s Δ ≈ 2.27. Surprisingly, the *E. histolytica* secretome exhibited a significantly lower mean disorder than its corresponding whole proteome (Table 3, Welch’s *t*-test, t ≈ −6.15, p < 0.000001). This difference was also associated with very large effect sizes (Cohen’s *d* ≈ −2.45, Hedges’ *g* ≈ −2.42, and Glass’s Δ ≈ −2.62). These data suggest that *E. histolytica* relies less on structurally flexible secreted proteins for host interaction and environmental adaptation than does *P. falciparum*.

**Table 3.** Descriptive statistics for mean per secretome and per proteome for *P. falciparum* and *E. histolytica*.

| Secretome vs Proteome Species<sup>a</sup> | n | Mean Disorder (±SD) | Welch's *t* | *p*-value | Cohen's *d* | Hedges' *g* | Glass's $\Delta$ | Effect Size Interpretation<sup>b</sup> |
| --- | --- | --- | --- | --- | --- | --- | --- | --- |
| Pf | 61 | 31.79(±26.40) | 3.07 | ~0.003 | 1.9 | 1.88 | 2.27 | Very large |
| Eh | 68 | 5.23(±9.98) | -6.15 | < 0.000001 | -2.45 | -2.42 | -2.62 | Extremely large |

<sup>a</sup>Species: *P. falciparum* 3D7 (Pf); *E. histolytica* (Eh)
<sup>b</sup>Effect size interpretation: negligible (<0.2), small (0.2–0.49), moderate (0.5–0.79), large (0.8–1.19), very large (1.2–1.99), and extremely large (≥2.0).

### Antigenicity of Disorder in *E. histolytica*

Historically, it was assumed that the high flexibility of intrinsically disordered regions makes them poor vaccine targets, as a lack of fixed structure may prevent the immune system from mounting a consistent and effective response [40,41]. More recently, this notion has been challenged. For example, in one study that examined immunodominant proteins of *Chlamydia* spp., protein disorder tendency was the best predictor of a candidate B-cell epitope [42]. Furthermore, several approved malaria vaccines (R21, RTS,S) target the circumsporozoite protein (CSP), a prominent surface protein of sporozoites. Interestingly, the disordered tetra-peptide repeats of CSP appear to be immunodominant and the targets of the protective antibodies that accumulate [43]. A second malaria vaccine targets the merozoite surface protein 2 (MSP2), which is predicted to be completely disordered [43]. Despite its disorder, the MSP2 antigen induced vigorous antibody responses and mediated protection in a Phase II trial in Papua New Guinea [43,44]. To explore if there is a relationship between protein disorder and antigenicity for *E. histolytica* proteins, we performed a secondary analysis of antigenic membrane proteins identified by Kumarasamy *et al*. [24]. The majority (8 out of 12) exhibited below-average intrinsic disorder when compared to the overall mean per protein disorder for the species (Table 4). Although we do not have access to a list of membrane proteins that are not antigenic, perhaps proteins with below-average disorder are immunodominant in *E. histolytica*.

**Table 4.** Average percent disorder of antigenic proteins of *E. histolytica*.

| Accession number<sup>a</sup> | Description | Average % Disorder |  |  | Mean % Disorder (±SD) | p<sup>b</sup> |
| --- | --- | --- | --- | --- | --- | --- |
|  |  | Disopred3 | IUPred3 | AUCpred |  |  |
| EHI_043640 | Actin, putative | 3.31 | 0.00 | 2.71 | 2.01±1.44 | 0.0046** |
| EHI_043010 | V-type ATPase, putative | 1.48 | 0.00 | 2.41 | 1.29±0.99 | 0.003** |
| EHI_012270 | Gal/GalNAc lectin heavy subunit | 4.59 | 0.00 | 6.22 | 3.60±2.63 | 0.0159* |
| EHI_044370 | Hypothetical protein, conserved | 2.06 | 0.00 | 1.91 | 1.32±0.94 | 0.0029** |
| EHI_130700 | Enolase, putative | 0.46 | 0.00 | 1.61 | 0.69±0.68 | 0.0022** |
| EHI_071590 | Protein disulfide isomerase, putative | 9.51 | 0.00 | 14.67 | 8.06±6.08 | 0.3032 |
| EHI_195760 | Alcohol dehydrogenase 3, putative | 0.00 | 0.00 | 1.04 | 0.35±0.49 | 0.0019** |
| EHI_004880 | Hypothetical protein | 6.53 | 0.00 | 4.08 | 3.54±2.69 | 0.016* |
| EHI_110350 | Thioredoxin, putative | 11.71 | 6.75 | 12.70 | 10.39±2.60 | 0.3712 |
| EHI_107290 | Actin, putative | 1.06 | 0.00 | 3.72 | 1.59±1.56 | 0.0042** |
| EHI_064680 | α-soluble NSF attachment protein, putative | 10.27 | 2.05 | 6.16 | 6.16±3.36 | 0.0643 |
| EHI_052860 | Heat shock protein 70, putative | 6.55 | 14.63 | 8.23 | 9.80±3.48 | 0.3383 |

<sup>a</sup> From Kumarasamy *et al.*, 2020
<sup>b</sup> Comparison to overall mean % disorder (per protein); calculated by Unpaired *t* test
\* $p < 0.05$ ; \*\* $p < 0.01$

Rahman *et al*. [42] reported that an effective approach for B-cell epitope prediction is to plot the disorder along protein sequences using the IUPred3 scale followed by antibody reactivity testing of peptides in the highest disordered regions. Therefore, we performed secondary analysis using data from two prior studies in which fragments of major vaccine targets were tested for their ability to block amoebic liver abscess formation in animal models of disease. One study focused on four fragments of the heavy subunit of the cell surface, galactose-inhibitable lectin (Hgl) [25], while the other investigated full-length and three fragments of the intermediate subunit of the cell surface, galactose-inhibitable lectin (Igl) [26]. The location of these fragments along the protein sequence, along with the predicted disorder profile are shown in Fig 4 (A,B). In both studies, protection against liver abscess served as the primary readout for vaccine efficacy. When compared to the overall mean per protein disorder for *E. histolytica,* full-length Hgl exhibited significantly lower intrinsic disorder than the whole proteome (3.60% ± 2.63%; *p*= 0.0164), while full-length Igl did not (7.84% ± 9.71; *p*= 0.4615). We plotted the average disorder of each antigen against its corresponding vaccine efficacy (Fig 4C,D). Interestingly, we observed strong correlations for both targets, albeit in opposite directions. Higher disorder was associated with increased protection for Hgl (Fig 4C) and decreased protection for Igl (Fig 4D). Other putative vaccine targets, such as the *Entamoeba histolytica* Surface Metalloprotease-1 (EhMSP-1) [45] and the Serine-Rich *Entamoeba histolytica* Protein (SREHP [46]), were not similarly analyzed. The EhMSP-1-based vaccine consisted of a mixture of different peptides, which precluded meaningful correlation with intrinsic disorder of a single polypeptide, and the SREHP study employed a plasmid DNA vaccine rather than a recombinant protein vaccine, limiting direct comparability.

**Fig 4.**
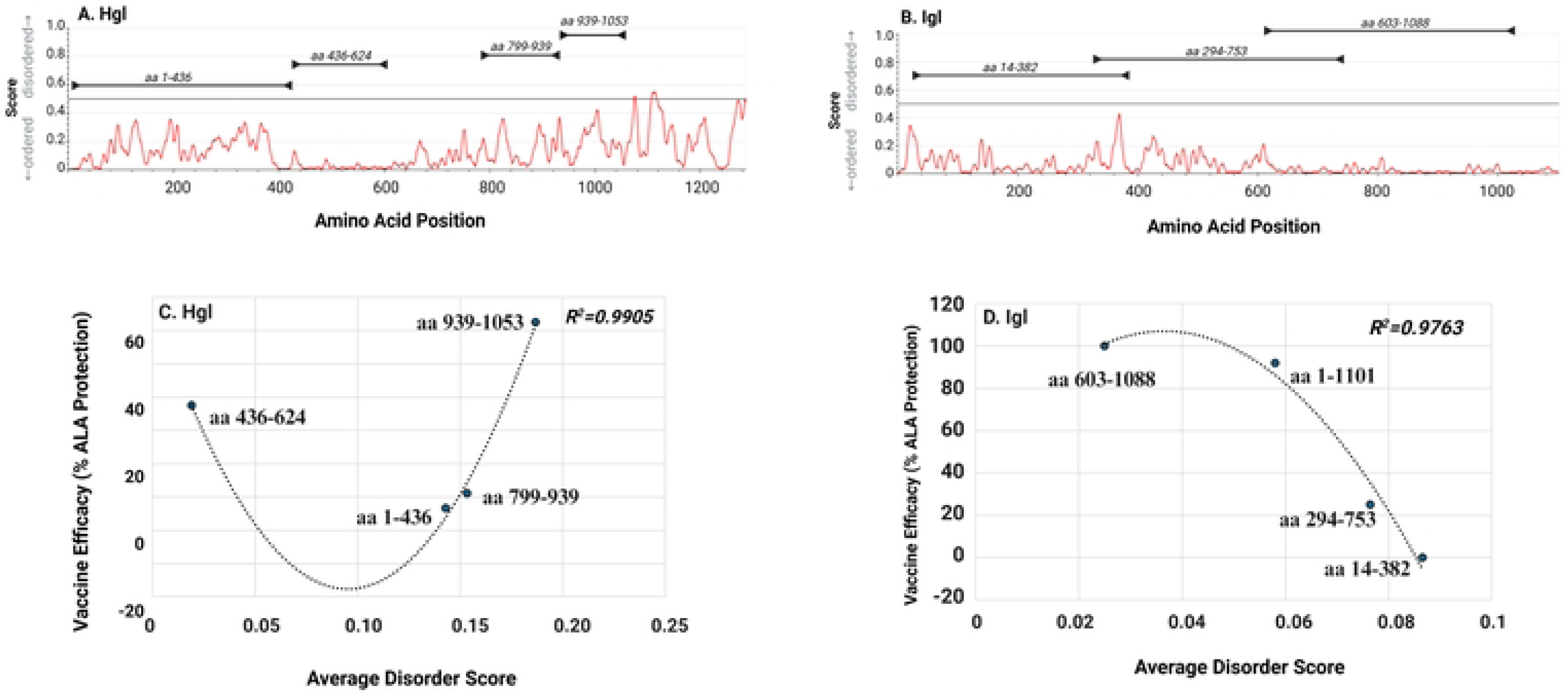
Relationship between intrinsic disorder and vaccine efficacy for Hgl and Igl antigens. (A–B) Predicted intrinsic disorder profiles across the amino acid sequences of Hgl (A) and Igl (B), with disorder scores plotted by residue position. Arrows indicate the boundaries of antigen fragments used for analysis. (C–D) Correlation between average disorder score of each fragment and vaccine efficacy (% ALA protection) for Hgl (C) and Igl (D). Individual data points correspond to specific fragments (amino acid ranges labeled), and dotted lines represent fitted regression curves (R² values shown). For Hgl, intermediate disorder regions exhibit lower efficacy, while highly disordered regions show increased protection. In contrast, for Igl, regions with lower to moderate disorder are associated with higher efficacy.

## Discussion

In this study we have undertaken the first proteome-wide analysis of intrinsic protein disorder in amoebozoans of the genus *Entamoeba*. We have shown that proteome-wide intrinsic disorder is significantly lower in the *Entamoebae* than in *P. falciparum* and *T. brucei*. Functional enrichment (Gene Ontology (GO)) analyses revealed a broadly conserved progression across all five organisms examined, in which protein functions shifted from metabolic and catalytic roles toward gene expression roles as intrinsic disorder increased. However, important organism-specific differences were observed. Notably, the *Entamoebae* maintained G-protein signaling across all levels of disorder, whereas *P. falciparum* maintained gene expression functions regardless of disorder level. In contrast, *T. brucei* was characterized by persistent enrichment of metabolic, transport, and redox processes across the disorder spectrum. Additionally, analyses of secretomes demonstrated that *E. histolytica* exhibited significantly lower disorder in its secreted proteins compared to its overall proteome, in contrast to *P. falciparum*, where secreted proteins were more disordered than its overall proteome. Finally, no clear correlation was observed between protein disorder and vaccine efficacy for *E. histolytica*.

In Eukarya, an evolutionarily conserved trend links low intrinsic disorder with metabolic and catalytic functions, and high intrinsic disorder with gene expression and regulatory roles [47,48]. However, this pattern represents a statistical trend and not an absolute functional dichotomy. For instance, some regulatory proteins may retain structured domains (e.g., DNA-binding motifs) [49], while some metabolic enzymes can contain disordered regions that contribute to regulation and allosteric interactions [50]. The five organisms studied here generally followed the evolutionary conserved trend.

Despite the shared trend, clear species-specific differences were observed. *P. falciparum* exhibited the strongest enrichment of transcriptional, translational, and RNA-processing functions across all levels of disorder. At first glance, this may be counterintuitive since the parasite’s gene expression program is tightly and temporally regulated, which might be expected to depend on a highly ordered machinery. For example, during the intraerythrocytic developmental cycle, *P. falciparum* undertakes host cell remodeling, including the export of proteins that alter erythrocyte structure and mechanics [51]. During schizogony, rapid rounds of DNA replication and cell division produce numerous merozoites, and invasion-related proteins are synthesized just prior to egress [52]. There is also strict control of antigenic variation through mutually exclusive expression of *var* genes [53]. These processes require precisely timed biosynthesis of proteins, all carried out by the parasite itself due to the lack of erythrocyte transcriptional capacity. The broad deployment of intrinsic disorder in *P. falciparum* gene expression categories suggests that while lower disorder might facilitate precision, higher disorder may be required for flexibility, which would enable transient interactions, assembly and disassembly of complexes, and responsiveness to changing intracellular conditions. Nevertheless, this interpretation remains speculative, and direct experimental studies will be required to determine the extent to which intrinsic disorder contributes to gene expression control in *P. falciparum*.

On the other hand, *T. brucei* retained enrichment for GO terms associated with enzymatic, redox, and transport processes even at higher disorder levels, indicating a broader functional deployment of IDPs. This may reflect its need for rapid metabolic adaptation in host environments. In the insect vector, *T. brucei* primarily relies on amino acid catabolism for fuel [54]. This shifts largely to a reliance on glycolysis in the mammalian host [55], which could, in turn, result in the accumulation of NADH, a potentially toxic byproduct of glycolysis. Thus, in the human host, the parasite must consistently fine-tune how much glucose it breaks down to manage its redox state, prevent the accumulation of toxic derivatives, and fuel continuous high energy immune evasion strategies. Immune evasion strategies are mediated through monoallelic expression and periodic switching of variant surface glycoprotein (VSG) genes [21], which requires extensive protein expression as well as membrane trafficking (both endocytic and exocytic). That enzymatic-, redox-, and transport-related GO processes were enriched, even in the higher disorder strata, suggests that intrinsic disorder is widely utilized to support metabolic plasticity and surface remodeling necessary for a successful host infection.

Strikingly, the *Entamoebae* uniquely maintained a high representation of signaling pathways, particularly those involving GTPases and phosphorylation, in the low, moderate, high and very high disorder categories. This was most pronounced for the two pathogenic species, *E. histolytica* and *E. invadens*. *E. histolytica* exhibits a remarkable expansion of small GTPases, including Rabs (>90) [56], alongside numerous Ras (∼10) and Rho (∼20) family members [57]. This far exceeds the repertoire of monomeric GTPases found in many other unicellular eukaryotes. This superfamily of GTP-binding proteins typically controls key processes such as vesicle trafficking, phagocytosis, actin cytoskeleton dynamics, and cell motility. In *E. histolytica*, all of these activities are essential for nutrient acquisition and host tissue invasion and are regulated by G-proteins. For instance, various isotypes of *E. histolytica* Rab7 and Rab11 regulate phagocytosis, trogocytosis (nibbling live cells), membrane recycling, and secretion [58–61]. A unique Rab, EhRabA, regulates phagocytosis and adhesion to host cells [62,63]. Rho family GTPases, regulate actin-mediated processes underlying amoeboid movement and phagocytic cup formation during ingestion of host cells [57]. *E. histolytica* must be able to perform all of these functions in the fluctuating environment of the host gut, which may include rapid changes in nutrient availability, microbiota composition, and host immune factors. Thus, a broad deployment of disorder in G-proteins likely enables swift adaptation.

Interestingly, there was a less pronounced enrichment of G-protein signaling GO terms in the highest disorder category, in the commensal, *E. dispar*. Since *E. dispar* does not aggressively phagocytose host tissue nor invade intestinal epithelial layers [19], it may not require an array of G-protein signaling function across all levels of intrinsic disorder. This suggests that the integration of GTPase signaling *at every level of protein disorder* is an evolutionary adaptation of pathogenic *Entamoeba* species.

A growing body of evidence, across diverse biological systems, supports the idea that secreted and host-interacting proteins are frequently enriched in IDRs. This trend is particularly well established in viruses, where many host-targeting proteins, such as HIV Tat [64] and adenoviral E1A [65], are highly disordered and use this flexibility to interact with multiple host factors and modulate transcriptional machinery. A similar principle extends to prokaryotes. Many bacterial type III secretion system effectors exploit intrinsic disorder because it facilitates translocation through secretion apparatuses and enables rapid folding and interaction within host cells. For example, *Shigella* Ipa invasion proteins contain extensive intrinsically disordered regions, while *Salmonella* SopE exhibits structural flexibility and low stability consistent with disorder [66,67].

In eukaryotic pathogens, this pattern is further exemplified in *P. falciparum*, in which exported proteins such as PfEMP1 (*var* gene product) and the knob-associated histidine rich protein (KAHRP) contain disordered regions that support adhesion, host cell remodeling, and immune evasion [9]. In *T. brucei* VSGs also exhibit highly flexible regions [68]. Collectively, these examples suggest that intrinsic disorder provides functional advantages for proteins operating at the host-pathogen interface, including conformational flexibility, binding promiscuity, and enhanced evolvability, making it a recurring and conserved feature of cell surface and secreted proteins throughout the tree of life.

Here, we demonstrate that the mean intrinsic disorder of the predicted secretome of *P. falciparum* was statistically higher than that of its whole proteome, which is consistent with the idea that secreted and host-interacting proteins are enriched in disorder. On the other hand, the mean disorder of the predicted secretome of *E. histolytic*a was statistically lower than that of its whole proteome. Given the principle outlined above, this finding was unexpected. It suggests that unlike other parasites, *E. histolytica* relies less on structurally flexible secreted proteins for host interaction and environmental adaptation. This unexpected structural bias in the *E. histolytica* secretome further illustrates that disorder is selectively deployed rather than universally favored, even in host-interacting systems.

There are several non-mutually exclusive explanations for the low disorder (high structure) pattern of the *E. histolytica* secretome. First, the *E. histolytica* secretome is enriched in pore forming and hydrolytic (e.g., cysteine proteases, glycosidases, phosphatases) proteins [69,70]. This repertoire is heavily biased toward macromolecular breakdown, which is linked to the parasite’s invasive, extracellular feeding strategy and tissue-destructive lifestyle. Thus, evolutionary selection in this genus may have favored a secretome with stable secondary and tertiary structures to support robust enzymatic activity. Second, inside of cells, proteins function in buffered, chaperone-rich environments. However, outside of cells, and particularly in the intestinal lumen, secreted proteins must function in an unpredictable environment that can vary in pH and ionic conditions, and possess digestive enzymes, host and bacterial proteases, mucous, and reactive oxygen and nitrogen species. Folded proteins (low disorder) may be better able to maintain activity under these fluctuating conditions because hydrophobic cores and other stabilizing interactions may preserve structure. Since IDRs are solvent exposed, flexible, and protease accessible, low disorder may also facilitate resistance to the intestinal environment by burying protease cleavage motifs. The functional demands imposed on secreted proteins of *E. histolytica* differ substantially from those imposed on secreted proteins of *P. falciparum*. Our data suggest that these demands may shape the balance between intrinsic order and disorder in secreted proteins.

Extensive efforts have been devoted to evaluating the heavy (Hgl) and intermediate (Igl) subunits of the Gal/GalNAc lectin as candidate vaccine antigens in *E. histolytica* [25,26]. The opposing relationship between intrinsic disorder and protection observed for these two antigens suggest that disorder is not a universal predictor of immunogenic efficacy in this species. Rather, different antigens may achieve protection through distinct mechanisms. For instance, a positive correlation between intrinsic disorder and protection, as was the case for Hgl, may reflect the capacity of disordered regions to present flexible, solvent-exposed epitopes, which are often enriched in linear B- and T-cell epitopes. In contrast, a negative correlation, where structurally ordered proteins are associated with stronger protection, as was the case for Igl, suggests that immunity may rely on conserved conformational epitopes, stable binding interfaces, or rigid domains. Although both studies employed the same inoculum sizes with a primary injection and two booster doses, we cannot exclude the possibility that the observed differences arise from variations in adjuvants, routes of administration (IP vs. IM), or animal models (hamsters vs. gerbils). Though it reinforces a need for a careful interpretation of disorder in vaccine design.

## Conclusion

Overall, our results support the notion that intrinsic disorder is not simply a passive structural feature, but a context-dependent property that is differentially integrated into cellular and host-parasite interaction networks across species. Thus, the variety of IDPs utilized in various contexts have diverged in accordance with an organism’s lifestyle and host interaction strategy. These findings highlight the role of intrinsic disorder in shaping parasitism and emphasize its importance in the evolution of diverse survival and virulence strategies of pathogens.

## Acknowledgements

This work was funded by National Institute of General Medical Sciences Grants: GM109094 (LAT) and GM146584. The funders had no role in the study design, data collection and analysis, decision to publish, or preparation of the manuscript. The authors thank Dr. Karl Franek (Bioscriptorium.com) and Dr. James Morris (Clemson University) for helpful discussions and critical evaluation of the manuscript.

## Supporting Information

**Supplementary File 1.** Excel file containing the complete protein datasets for all five species analyzed. For each protein, the table includes accession number, protein description, and predicted intrinsic disorder scores obtained using Disopred3 [27], IUPred3 [28], and AUCpreD [29]. Functional annotation is provided through associated Gene Ontology (GO) terms for Cellular Component, Molecular Function, and Biological Process. Enzyme Commission (EC) numbers are included where applicable.

**Supplementary File 2.** Excel file containing the top 10 enriched Gene Ontology (GO) terms for Cellular Component, Molecular Function, and Biological Process categories for each of the five species analyzed. Enrichment results are stratified by predicted intrinsic disorder levels of proteins (0–10%, 10–30%, 30–70%, or >70%), with terms ranked by enrichment significance within each disorder bin for each species.

## Notes

### Competing Interest Statement

The authors have declared no competing interest.

